# A human *ex vivo* dengue virus neutralization assay identifies priority antibodies and epitopes for vaccines and therapeutics

**DOI:** 10.1101/516195

**Authors:** Trung Tuan Vu, Hannah Clapham, Van Thi Thuy Huynh, Long Vo Thi, Dui Le Thi, Nhu Tuyet Vu, Giang Thi Nguyen, Trang Thi Xuan Huynh, Kien Thi Hue Duong, Vi Thuy Tran, Huy Le Anh Huynh, Duyen Thi Le Huynh, Thuy Le Phuong Huynh, Thuy Thi Van Nguyen, Nguyet Minh Nguyen, Tai Thi Hue Luong, Nguyen Thanh Phong, Chau Van Vinh Nguyen, Gerald Gough, Bridget Wills, Lauren B. Carrington, Cameron P. Simmons

## Abstract

**Background:** Dengue is the most prevalent arboviral disease, for which neither effective vaccines nor antivirals are available. Clinical trials with Dengvaxia, the first licensed dengue vaccine, show the conventional in vitro plaque reduction neutralization test (PRNT) failed to discriminate between neutralizing and non-neutralizing antibodies. A number of human monoclonal antibodies (mAbs) were characterized by PRNT as being neutralizers of virus infectivity for mammalian cells.

**Methodolody/Principle findings:** We developed a neutralization assay and tested the capacity of 12 mAbs to neutralize the infectiousness of dengue patient viremic blood in mosquitoes. We identified minimum concentrations of a subset of mAbs required to achieve dengue virus neutralization, and modelled the impact of a therapeutic mAb candidate on viremia.

Five of the 12 mAbs (14c10, 2D22, 1L12, 747(4)B7, 753(3)C10), all of which target quaternary epitopes, potently inhibited dengue virus infection of *Ae. aegypti*. The potency of several mAbs was compromised in the context of patients with secondary serological profiles, possibly reflecting competition between the exogenously-added mAbs and the patient’s own antibody responses at or near the target epitopes. The minimum concentrations that mAbs neutralized DENV ranged from 0.1 – 5 µg/mL. An Fc-disabled variant of mAb (14c10-LALA) was as potent as its parent mAb. Within-host mathematical modelling suggests infusion of 14c10-LALA could bring about rapid acceleration of viremia resolution in a typical patient.

**Conclusions/Significance:** These data delivered a unique assessment of anti-viral potency of a panel of human mAbs. Results support the advancement of dengue virus neutralization assays, and the development of therapeutics against flaviviruses, to which dengue virus and Zika virus belong.

**Author summary:** Dengue is the most prevalent arboviral disease affecting humans. There are no therapeutics for the disease. Antibody-mediated immunity against dengue is also not well-understood, as shown by the failure of the conventional neutralization assay used to predict the efficacy of Dengvaxia, the first licensed vaccine for the disease. It is likely that the neutralization assay targets non-neutralizing antibodies, but there are no validation assays available. To this end, we developed a novel virus neutralization assay, employing *Aedes aegypti* mosquitoes and viremic blood from dengue patients, to examine the virus-neutralizing potency of 12 human-derived monoclonal antibodies (mAbs). While all of these mAbs neutralized dengue virus using the conventional assay, seven of them failed to block dengue virus infections of mosquitoes using our assay. The remaining five mAbs neutralized at least one serotype of dengue virus and the minimum neutralizing concentrations of range from 0.1 – 5 µg/mL. Using the minimum neutralizing concentration of a therapeutic mAb candidate, we investigated the impact of the mAb on viremia using a mathematical model and found the mAb accelerated the reduction of viremia. The results support the advancement of dengue virus neutralization assays, and the development of therapeutics for dengue.

## Introduction

Dengue virus (DENV) infections are highly prevalent in the tropical and subtropical world (1). Following a primary DENV infection it is widely accepted that an individual develops long-lived clinical immunity to the infecting serotype but not to other serotypes. DENV infection can be subclinical, or result in a febrile syndrome that in a small percentage of patients is complicated by vascular leakage, thrombocytopenia and altered hemostasis, usually between the fourth and sixth days of illness (2). The risk of clinically important complications is higher when an individual is infected for a second time with a different DENV serotype from the first (3). In this situation, cross-reactive antibodies are hypothesized to enhance the viral infection (4).

Antibodies are central to the concepts of dengue pathogenesis and naturally-acquired or vaccine-elicited immunity. For example, the induction of virus neutralizing antibodies by vaccination is the goal of all candidate dengue vaccines in clinical development (5). The only-licensed dengue vaccine, Dengvaxia, was developed on the basis that it could elicit antibodies that neutralized all four serotypes of DENV *in vitro* (6), but the vaccine mediated an increased risk of dengue in vaccine recipients who were seronegative at baseline (7).

Despite the agreed centrality of antibodies to dengue pathogenesis, laboratory correlates of immunity (or disease enhancement risk) have not been tightly defined or standardized. A general consensus is higher serum concentrations of neutralizing antibodies are associated with reduced risk of dengue (8, 9), but results from plaque reduction neutralization test (PRNT), used to quantify neutralizing antibody responses, failed to predict vaccine efficacy in Dengvaxia clinical trials. A possible explanation is that PRNT relies on referenced DENV strains and mammalian cells which do not mimic natural DENV infection. As such, the availability of evidence-based and agreed correlates of immunity (or disease risk) could help fast track development of new dengue vaccines with better efficacy and safety profiles than Dengvaxia (10). This goal may require “second generation” virus neutralization assays that improve upon the predictive value of PRNT.

The ability to generate and characterize human monoclonal antibodies (mAbs) from dengue immune donors has provided insights into the type of antibodies that might be desirable to elicit via immunization, or alternatively, to use as therapeutic agents. Most prominent are those human mAbs identified as being potent neutralizers of virus infectivity for mammalian cells (Table S1) (11-17). Many of these neutralizing mAbs bind to viral envelope proteins that have quaternary structures (18-21).

Here we developed a <u>vi</u>remic <u>b</u>lood <u>n</u>eutralization <u>a</u>ssay (ViBNA), in which blood from DENV-confirmed patients was spiked with mAbs, and was then used to feed *Aedes aegypti* mosquitoes. We use this assay to characterize neutralization capacity of a panel of 12 anti-DENV human mAbs with different serotype specificities and epitope binding characteristics. We identified a short-list of mAbs with their minimum concentrations that neutralized DENV using ViBNA. Further, we modelled the anti-viral effect of a therapeutic LALA mAb dosing to deliver predictions that could influence the design of future clinical trials. These results inform dengue therapeutic and the advancement of a new generation of immune correlate assays for vaccine development.

## Methods

### Viremic Blood Neutralization Assay (ViBNA)

In ViBNA, viremic blood was drawn directly from acutely-infected dengue patients, and the infection of blood-fed mosquitoes was the assay endpoint to measure neutralization characteristics and potency of monoclonal antibodies (mAbs).

Viremic blood was drawn from NS1-positive dengue patients and spiked with various concentrations of individual mAbs. The blood-antibody mixture was incubated at 37°C for 30 minutes for virus neutralization to occur. The 30-minute duration is a standard time for mAbs to neutralize virus, as with other neutralization assays. Positive and negative controls were prepared in parallel. The blood-antibody mixture was then maintained at 37°C in artificial membrane feeders, attached to a circulating warm water system. Mosquitoes were allowed to feed on the antibody-blood mixture for maximum 1 hr. Between 20-50 wild-type *Aedes aegypti* were included for each preparation and engorged mosquitoes were maintained for 5-7 days, allowing non-neutralized dengue virus (DENV) to amplify in the mosquitoes. Those surviving the incubation period were sacrificed, homogenized, and tested for the presence of DENV RNA using RT-PCR procedures described elsewhere (22).

Clone names and characteristics of the 12 human mAbs are described in Table S1. The mAb 14c10 was obtained with two separate variants, wildtype and LALA. The LALA modification abrogates mAb binding to Fc receptors.

The positive control was hyper-immune plasma, created by pooling convalescent plasma samples of 119 dengue-confirmed patients with any of the four serotypes. The plasma pool was then heated at 56°C for 30 minutes to inactivate complement. Negative control was 0.9% saline. Both controls were diluted 1:9 in viremic blood before mosquito exposures.

### Dengue patient cohorts and diagnostic tests

Eligible patients; who were admitted to inpatient wards at the Hospital for Tropical Diseases, HCMC, were enrolled between April 2013 and July 2017. The inclusion criteria were: age ≥15 years; <96 hours of fever history; and a positive NS1 rapid test to confirm DENV infection. There were no exclusion criteria. A single 3-5 mL venous blood sample was drawn from each participant on the day of enrolment, or occasionally, the following day, for mosquito feeding. Subsequently RT-PCR was performed to quantify viremia levels following established methodology (23). Panbio’s Indirect IgG ELISA was also employed to examine the presence of DENV plasma IgG. The blood samples that underwent RT-PCR and DENV-reactive IgG test were separate aliquots from those used to feed mosquitoes.

### Ethics statement

This study was approved by the Ethics Committee of Hospital for Tropical Diseases, Vietnam (CS/ND/12/16, CS/ND/16/27) and the Oxford University Tropical Research Ethics Committee, UK (OxTREC 30-12, OxTREC 45-16). All patients provided written informed consent.

### *Aedes aegypti* mosquitoes

We used 3-5 days-old F3 laboratory-reared *Aedes aegypti* mosquitoes in these experiments. All mosquitoes were derived from field-sourced materials (eggs or larvae) collected in HCMC. Mosquitoes were maintained at 28°C, with a 12:12 light:dark cycle and 80% relative humidity. Mosquito colonies were blood-fed on healthy consent humans and had access to 10% sucrose *ad libitum*.

### Data screening and statistical analysis

We excluded blood meals from the analysis for any of the following reasons: 1) non-infectious, i.e. mosquitoes in the negative control were DENV negative; 2) all mosquitoes of either negative control or positive control died before harvesting; 3) the number of harvested mosquitoes from a single cohort was fewer than four. Due to the nested nature of the data, i.e. mosquitoes fed on each preparation of blood of the same dengue patient, a marginal logistic regression model was used to evaluate the neutralization capacity of any given mAb, relative to the negative control from within each antibody treatment. The model was multivariable, adjusting for the covariate of plasma viremia. Neutralizing mAbs were classed as those that generated odds ratio values less than 0.1. A *p* value of less than 0.001, adjusted for multiple-comparisons by the Bonferroni correction method, was statistically significant. To compare the percentage reduction between DENV-specific IgG-positive and -negative blood meals, we used the Wilcoxon rank sum test.

The percentage reduction in mosquito infections was calculated for each feed as follows: 100% - 100% x (the proportion of DENV infected mosquitoes in the mAb group / the proportion of DENV-infected mosquitoes in the saline group).

### *In vivo* modelling of a therapeutic mAb application

We modelled the impact of 14c10-LALA on DENV-1 viremia with the minimum concentration required to neutralize DENV infection. The clinical dosing of the mAb was estimated for intravenous infusion, with assumptions of an average human bodyweight of 70 kg and the amount of body plasma of 5.5 L.

We assessed the possible impact of 14c10-LALA to dengue viremia kinetics, by extending a previously published model of dengue viral replication (24, 25) to include different efficacies of 14c10-LALA. We simulated the model using samples from the posteriors of the model fits from this previous work. The action of the mAb was simulated as an increase in the rate at which virus was cleared. We assumed the action of the mAb, administered once, did not change over time and not affect the growth or the binding of patients’ antibodies.

Two scenarios were simulated; high and low clearance by 14c10-LALA, corresponding to a viral clearance rate of 10,000 and 100. We simulated the mAb as being administered at day 9 after infection, corresponding to on average day 3 of symptoms (26), the earliest day patients usually report to hospital.

## Results

### Neutralization capacity of monoclonal antibodies (mAbs)

In our patient population, DENV-1 and DENV-4 were the most prevalent serotypes. The median plasma DENV-RNAemia level varied by serotype and was highest for DENV-1 (Fig S1).

### Least potent mAbs

At a concentration of 10 μg/mL; 1C19, 1M7, 22.3, 82.11, 87.1, 5J7 and 1F4 were the least potent at neutralizing the infectivity of DENV viremic blood for *Aedes aegypti* mosquitoes (Fig 1 and Table S2). Only 1F4 (a mAb previously characterized as DENV-1 specific) provided partial blocking of DENV-1 infection of mosquitoes (OR 0.18, 95%CI 0.10 – 0.34, P < 0.001).

**Fig 1.**
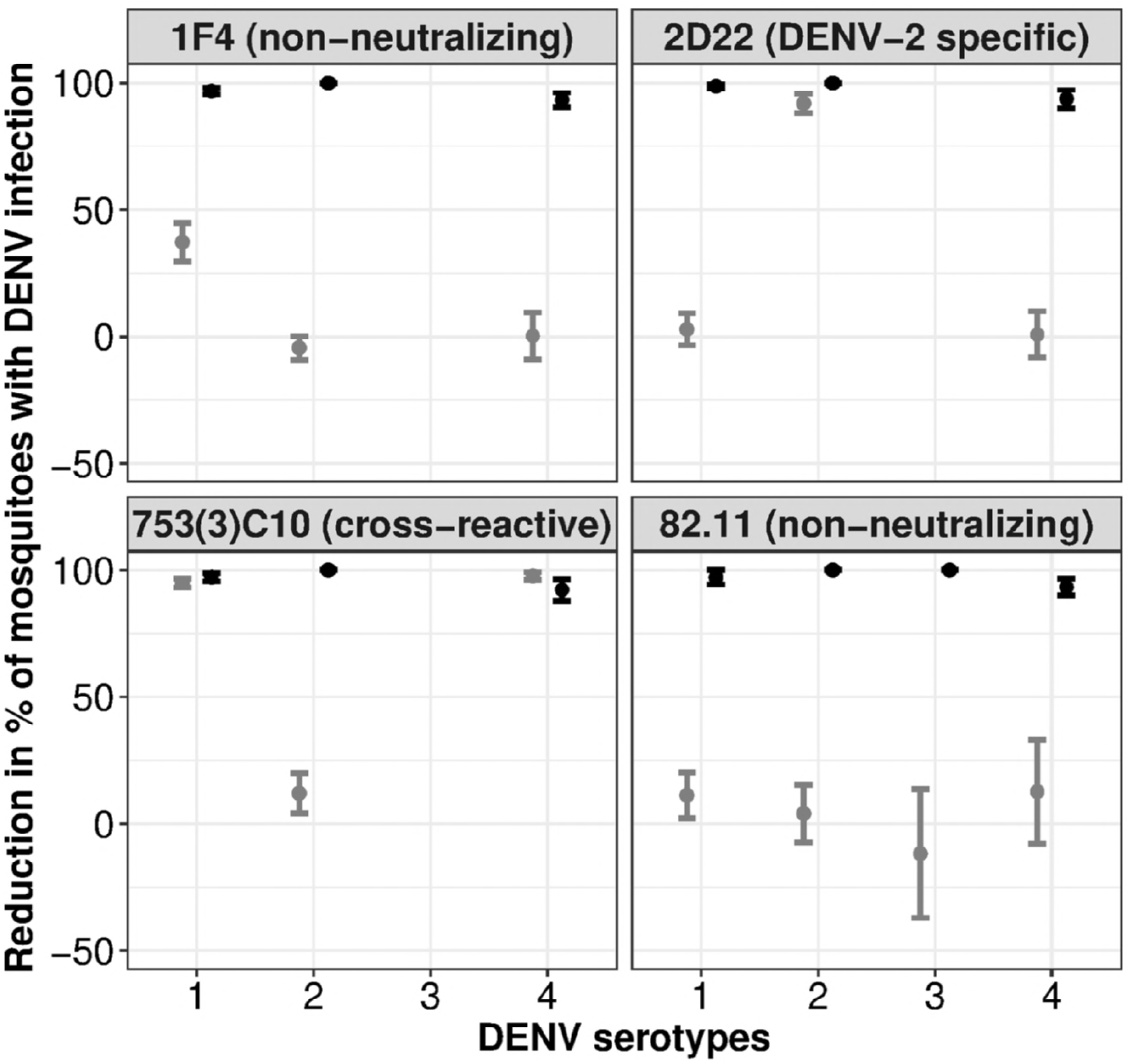
An example of neutralizing and non-neutralizing monoclonal antibodies (mAbs), according to the ViBNA. At the top of each panel are the mAb clone names. While 1F4 and 82.11 failed to neutralize DENV of any serotypes, 2D22 is DENV-2 specific and 753(3)C10 is cross-reactive against DENV-1 and DENV-4. This is indicated by the y-axis values of 100, meaning the mAb neutralize DENV, relative to the negative control. Means and standard errors of test results are calculated from three or more independent ViBNA measurements. Positive control is in black while tested mAbs are in gray.

### Most potent mAbs

Five mAbs, all of them recognizing quaternary epitopes on the virion surface, reproducibly neutralized DENV in viremic blood (Fig 1 and Table S2). The mAbs 14c10 (DENV-1 specific), 1L12 and 2D22 (both DENV-2 specific) blocked viral infection of mosquitoes at 10 µg/mL. The E-dimer epitope (EDE)-specific mAbs 747(4)B7 and 753(3)C10, both previously nominated as serotype cross-reactive, had different potency characteristics at the highest concentrations tested (3.7 µg/mL and 5 µg/mL, respectively). 753(3)C10 blocked DENV-1 and DENV-4 but did not prevent mosquitoes from acquiring DENV-2 infection. 747(4)B7 was also highly potent against DENV-1, but only partially blocked DENV-2 and DENV-4. There were insufficient blood meals to characterize the potency of any of these mAbs against DENV-3.

We hypothesized that the neutralization capacity of the serotype cross-reactive mAbs 747(4)B7 and 753(3)C10 might be influenced by the presence of pre-existing patient-derived antibodies, such as those binding to the E fusion loop region that compete and block access to the EDE quaternary epitope. We therefore tested for the presence of patient-derived IgG to the DENV virion (assessed by Panbio IgG indirect ELISA) in viremic blood samples. Consistent with the competition hypothesis, we found that the potency of 747(4)B7 was greatest in viremic blood samples devoid of measurable pre-existing IgG to the DENV (Wilcoxon rank sum test, p=0.023). A similar pattern was observed with 753(3)C10 but this was not statistically significant. In contrast, the potency of the DENV-1 specific mAb 14c10 was unaffected by pre-existing DENV-reactive IgG (Fig 2), suggesting that epitope-binding competition with other antibodies was less relevant to serotype-specific mAbs.

**Fig 2.**
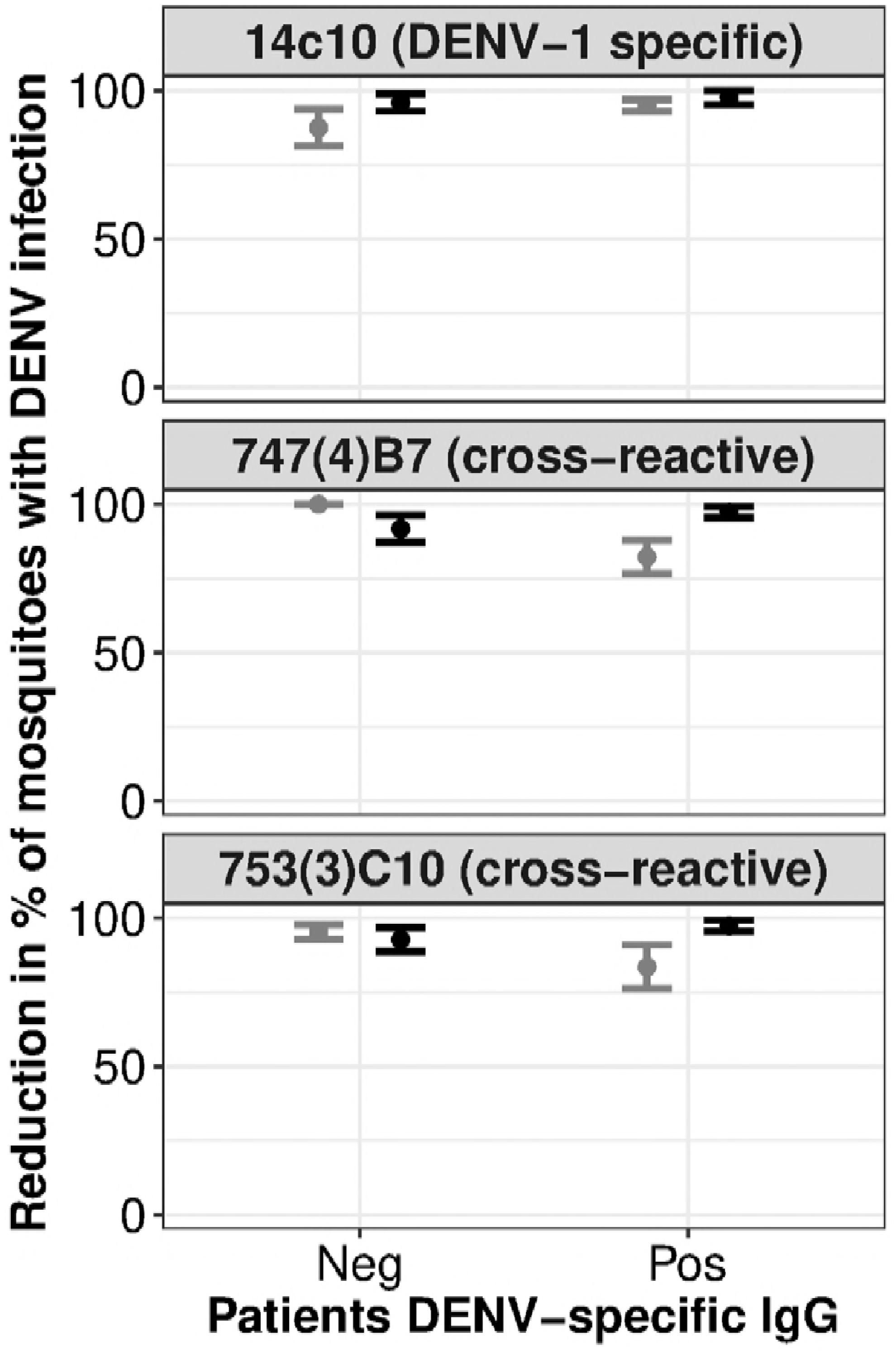
Reduced neutralization capacity of cross-reactive monoclonal antibodies (mAbs) in IgG-positive dengue patients. At the top of each panel are the mAb clone names, and the neutralization characteristic of that mAb, according to ViBNA. The y-axis value of 100 indicates the mAb neutralizes DENV, relative to the negative control. Due to the serotype specificity, only results of DENV-1 blood meals are shown with 14c10. Means and standard errors of more than two replicates are shown. Positive control is in black while tested mAbs are in gray.

Collectively, these experiments identified several mAbs capable of dramatically and quickly reducing the titer of infectious virus in patients’ blood to below the concentration required to infect *Aedes aegypti* mosquitoes.

### Minimum inhibitory concentrations of a subset of neutralizing mAbs

We investigated the lowest concentration of 14c10, 747(4)B7 and 753(3)C10; required to neutralize DENV in viremic blood. 14c10 neutralized DENV-1 at 10, 1 and 0.1 µg/mL but not at 0.01 µg/mL. The minimum concentration of 14c10 was also independent of Fc receptor engagement, as the Fc disabled variant 14c10-LALA was just as potent as the parent mAb (Fig 3). DENV-1 infection was completely inhibited in mosquitoes by 747(4)B7 at 3.7 µg/mL and 0.37 µg/mL, but not at lower doses, and the mAb generally failed to block DENV-4 infection at all concentrations tested (Fig S2). 753(3)C10 was less potent than 747(4)B7 against DENV-1, blocking only at the highest concentration tested, 5 µg/mL. 753(3)C10 also blocked the infectivity of DENV-4 viremic blood at 5 µg/mL, but failed to neutralize virus when diluted any further (Fig S3).

**Fig 3.**
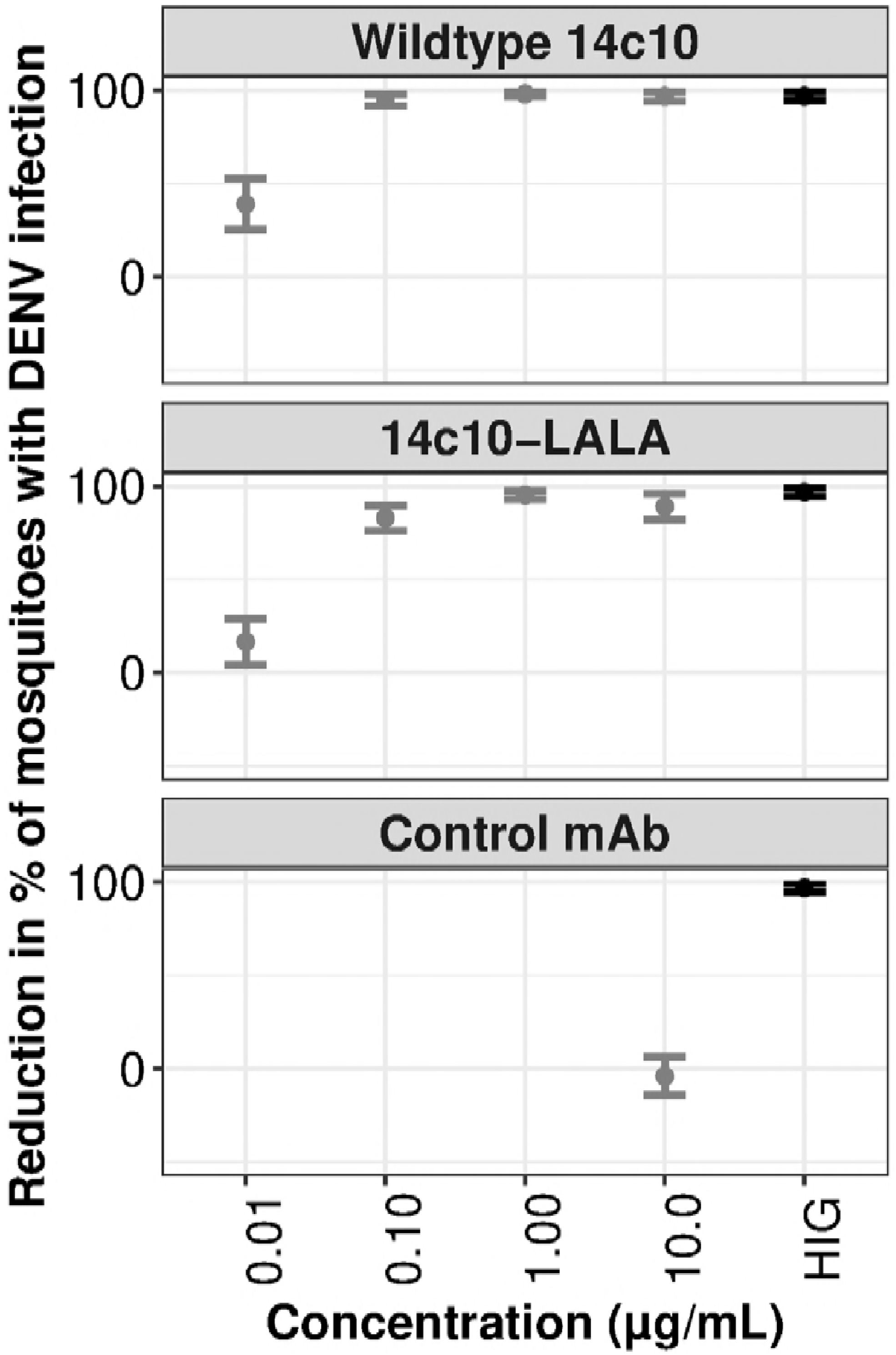
Both 14c10 variants neutralize dengue virus (DENV) at concentrations as low as 0.1 µg/mL. The y-axis value of 100 means mAb blocks DENV infection of mosquitoes completely, relative to the negative control. The isotype control mAb binds to respiratory syncytial virus. Means and standard errors of more than two replicates are shown. Positive control is in black while negative control is in gray.

### Modelled outcomes of therapeutic application of a potent, virus neutralizing mAb on DENV viremia

In a simplistic model, a blood concentration of 0.1 µg/mL mAb could be achieved in a 70 kg person with 5.5 L of blood, by intravenous infusion of 7 µg of mAb per kg bodyweight. For the majority of individuals at the 10,000-fold accelerated clearance, viremia immediately fell and continued to decline to undetectable levels 3-5 days before viremia would have been resolved naturally (Fig 4). A few individuals however observed a two-phase decline. At 100-fold, an immediate decline happened for many individuals, however a larger proportion of individuals saw a two-phase decline (Fig S4). This two-phase decline was observed when, at the point of mAb administration, the total antibody, comprising patient’s antibody plus 14c10-LALA was not yet sufficiently high to cause viral titers to start declining immediately. This happened more often for individuals experiencing a primary infection where natural antibody titers were in general lower at the time of administration than secondary individuals. However for all individuals, viremia decline started soon after administration. These modelling data suggest potent virus neutralizing mAbs could plausibly bring about clinically relevant reductions in the magnitude and duration of DENV viremia.

**Fig 4.**
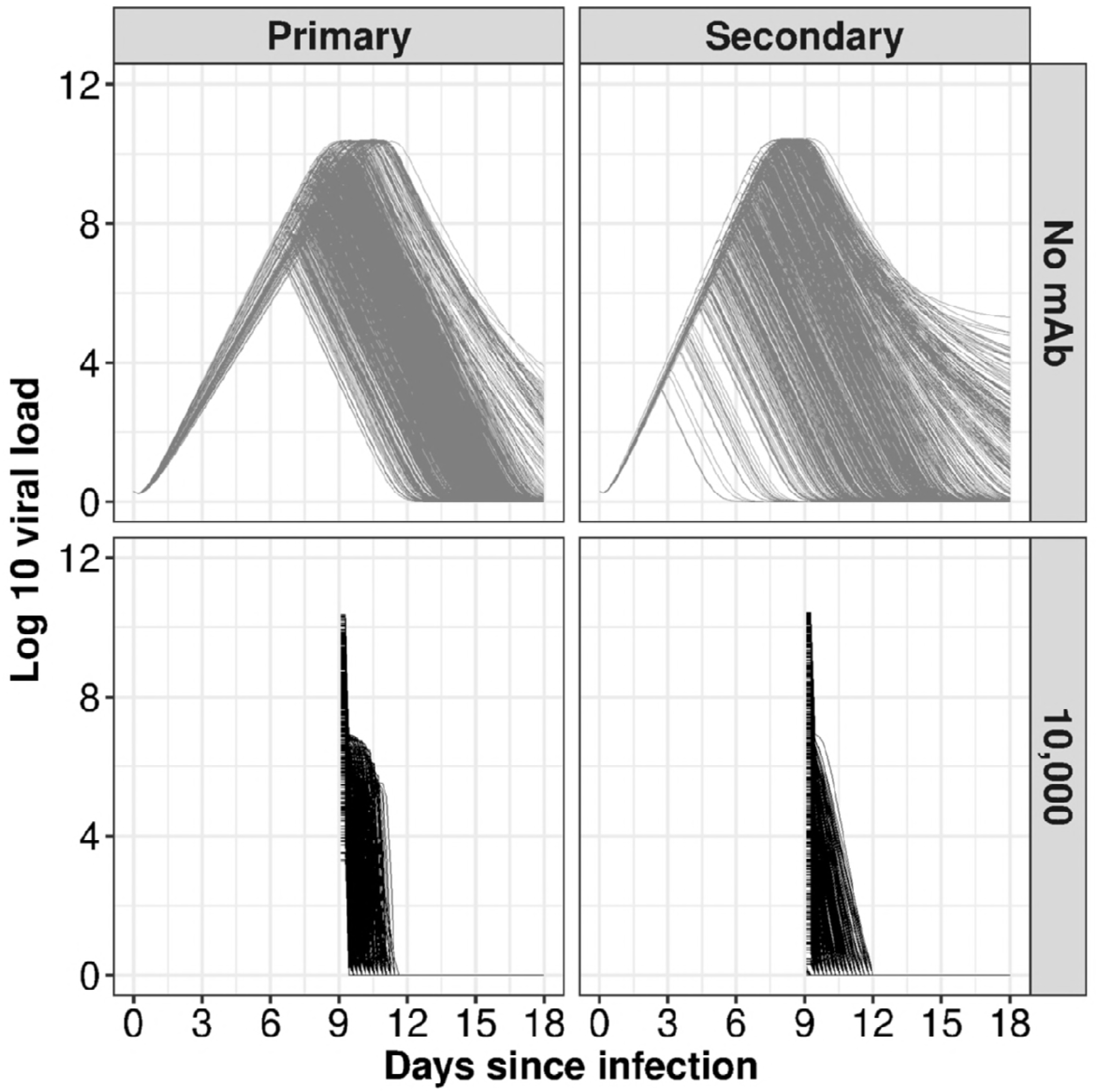
Modelled outcomes of administering 14c10-LALA on dengue virus-1 viremia. Log10 viremia is on y-axis and days since infection is on x-axis. Gray lines show the viral titers, described elsewhere (24), without monoclonal antibodies. Accelerated clearance rates of 10,000-fold are in black. Serological status (primary vs. secondary dengue infection in enrolled patients) is shown in the columns.

## Discussion

Here we characterize the neutralization capacity of a panel of well-characterized human monoclonal antibodies (mAbs) to block the infectivity of human viremic blood for *Aedes aegypti* mosquitoes. The mAbs that recognized quaternary epitopes on the virion surface were the most potent. The potency of the EDE specific mAbs 747(4)B7 and 753(3)C10 were compromised in the context of viremic blood from patients with secondary serological profiles, possibly because of competition at or around the epitope binding site by patient-derived anti-dengue virus (DENV) IgG. An Fc-disabled variant of the DENV-1 specific mAb 14c10 was as potent as its wild-type parent, confirming that Fc receptor engagement was unnecessary for virus neutralization. Within-host modelling suggests clinical infusion of a potent mAb such as 14c10-LALA could bring about rapid acceleration of viremia resolution.

For decades, the research community has been relying upon the PRNT to measure DENV-neutralizing antibodies after vaccination or natural infection. Coupled to this has been a decades-old working assumption that the PRNT (or related assays) accurately measure a correlate of clinical immunity. The validity of this assumption has been challenged by outcomes in trials of the only licensed dengue vaccine (Dengvaxia), where vaccine recipients who seroconverted in the PRNT were not always clinically immune to natural infection (6). This serves as the motivating example for why more accurate immune markers of clinical immunity to dengue must be identified. A limitation of PRNT is the use of referenced DENV strains and mammalian cells, irrelevant to natural DENV infections. In contrast, a key strength of our viremic blood neutralization assay (ViBNA) is it enables testing of mAbs in the physiologically relevant environment of freshly-collected, viremic blood from acute dengue patients. As such, ViBNA identified mAbs that potently neutralized virus in a “natural history” context. Were such antibodies elicited in sufficient concentrations by vaccination, it would be reasonable to hypothesize that they could mediate immunity to clinical disease, or potentially even mediate sterilizing immunity.

The magnitude of virus neutralization mediated by some mAbs was striking. For example, in acute viremic blood samples that contained >1,000 times the mosquito infectious dose of DENV-1 (Fig S1) (27), the mAbs very rapidly blocked the blood’s infectiousness. As expected, this occurred independently of Fc receptor engagement. The mAbs 747(4)B7 (EDE2) and 753(3)C10 (EDE1) performed similarly, but with the caveat that their potency was lower in blood samples from secondary dengue cases. Of relevance, both 747(4)B7 and 753(3)C10 mAbs target the same residues from three polypeptide segments on domain II of the envelope dimer: the *b* strand (amino acids 67–74, bearing the N67 glycan), the fusion loop and residues immediately upstream (amino acids 97–106), and the *ij* loop (amino acids 246–249). Thus we hypothesize that the “endogenous” IgG response in these patients (particularly antibodies binding to the fusion loop, a long-established target of serotype cross-reactive but poorly neutralizing antibodies (11, 16)), sterically interferes with 747(4)B7 and 753(3)C10 binding their epitopes. The implication is therapeutic approaches using EDE1 or EDE2 mAbs against DENV (16) or Zika virus (28) may have greater efficacy in cases with primary rather than secondary flavivirus infections. Finally, our “application” of a potent therapeutic mAb to a previously described within-host model of DENV infection in humans (29) suggests significant reductions in the course of viremia could be achieved with highly potent mAbs. The challenge is to provide proof of concept that reductions in the duration and magnitude of viremia can deliver clinically useful reductions in the duration and severity of illness.

What of antibodies that were not potent in the ViBNA? We cannot conclude that these mAbs lacked virus neutralization activity, only that they lacked sufficient potency to reduce the titer of infectious virus in the patients’ blood samples (against which they were tested) to below the mosquito infectious dose. A common feature of mAbs that had no or intermediate potency in the ViBNA was that they recognized epitopes available on the soluble envelope dimer (e.g. 22.3 and 1C19). This, and other work (21), suggests that vaccine approaches that present intact virions to the immune system, e.g. live attenuated viruses (30) or inactivated whole virion vaccines, are most likely to elicit the kind of quaternary-epitope-specific antibodies that in ViBNA were most potent.

Our study had several limitations. First, the amount of DENV varied between patients, which may influence the binding activity of mAbs, and the infection of mosquitoes. We could not normalize plasma viremia because the infectivity of DENV in viremic blood will be reduced after additional experimental procedures. Alternatively, we employed the marginal logistic regression model to control for the confounding factor of viremia. Second, we could not score the potency of all mAbs against all four serotypes of DENV because this was governed by the serotype prevalence. Third, the ViBNA effectively only detected virus neutralization when the mAb could reduce the infectious titer to below the mosquito infectious dose. Thus, rather than generating an end-point titer, the assay identifies a minimum concentration of mAb capable of consistently neutralizing virus infectivity to mosquitoes across a range of patient blood samples. Despite these limitations, this study has delivered a unique measurement of anti-viral activity for a panel of human mAbs and thereby could assist therapeutic and vaccine development. This is especially pertinent when the complexity and risks associated with dengue vaccine development have escalated, rather than diminished, since the first vaccine (Dengvaxia) was licensed.

## Acknowledgments

We thank collaborators for providing monoclonal antibodies (mAbs): Paul A. MacAry, GlaxoSmithKline for 14c10 and 14c10-LALA; Aravinda de Silva for 2D22, 1L12, 1F4, 1M7, 1C19, and 5J7; Antonio Lanzavecchia for 22.3, 82.11, and 87.1; Gavin Screaton for 747(4)B7 and 753(3)C10. We appreciate the contribution of patients and their families, and that of staff in OUCRU, and in the Hospital for Tropical Diseases.

## Supporting information Captions

**Fig S1.** Flow chart of the processing of viremic blood neutralization assay (ViBNA). Briefly, groups of mosquitoes fed on viremic blood spiked with the monoclonal antibodies. Engorged mosquitoes were then collected and tested for dengue virus (DENV) RNA. Excluded data originated from cases of non-infectious blood meals or from cases where all mosquitoes within the positive or negative control cohorts died before being collected, and therefore could not be assessed. Viremic blood samples were independently assessed for DENV serotype and viremia.

**Fig S2.** 747(4)B7 neutralized DENV-1 at a concentration as low as 0.37 µg/mL. Means and standard errors of more than two replicates are shown. Hyper-immune dengue virus (DENV)-reactive globulin (HIG), used as positive control, is in black while monoclonal antibodies are in gray. The y-axis value of 100 means that the mAb blocks DENV infection of mosquitoes completely, relative to the negative control.

**Fig S3.** 753(3)C10 neutralized DENV-1 at a concentration as low as 5 µg/mL. Means and standard errors of more than two replicates are shown. Hyper-immune dengue virus (DENV)-reactive globulin (HIG), used as positive control, is in black while monoclonal antibodies are in gray. The y-axis value of 100 means the mAb blocks DENV infection of mosquitoes completely, relative to the negative control.

**Fig S4.** Modelled outcomes of administering 14c10-LALA on dengue virus-1 viremia. Log10 viremia is on y-axis and days since infection is on x-axis. Gray lines show the viral titers, described elsewhere (24), without monoclonal antibodies. Accelerated clearance rates of 100-fold are in black after a single administration of 7 µg/kg of 14c10-LALA at day 9 of infection. Serological statuses are arranged by columns.

**Table S1.** Previously characterized properties of tested monoclonal antibodies (mAbs)

**Table S2.** Neutralization capacity of all monoclonal antibodies (mAbs) in the viremic blood neutralization assay (ViBNA)

